# Virtual reality validation of naturalistic modulation strategies to counteract fading in retinal stimulation

**DOI:** 10.1101/2021.11.17.468930

**Authors:** Jacob Thomas Thorn, Naïg Aurelia Ludmilla Chenais, Sandrine Hinrichs, Marion Chatelain, Diego Ghezzi

## Abstract

**Objective:** Temporal resolution is a key challenge in artificial vision. Several prosthetic approaches are limited by the perceptual fading of evoked phosphenes upon repeated stimulation from the same electrode. Therefore, implanted patients are forced to perform active scanning, via head movements, to refresh the visual field viewed by the camera. However, active scanning is a draining task, and it is crucial to find compensatory strategies to reduce it.

**Approach:** To address this question, we implemented perceptual fading in simulated prosthetic vision using virtual reality. Then, we quantified the effect of fading on two indicators: the time to complete a reading task and the head rotation during the task. We also tested if stimulation strategies previously proposed to increase the persistence of responses in retinal ganglion cells to electrical stimulation could improve these indicators.

**Main results:** This study shows that stimulation strategies based on interrupted pulse trains and randomisation of the pulse duration allows significant reduction of both the time to complete the task and the head rotation during the task.

**Significance:** The stimulation strategy used in retinal implants is crucial to counteract perceptual fading and to reduce active head scanning during prosthetic vision. In turn, less active scanning might improve the patient’s comfort in artificial vision.

## 1. INTRODUCTION

Retinal prostheses provide artificial vision to blind people affected by photoreceptor degeneration through the electrical stimulation of the retina, inducing the perception of localised bright spots of light, called phosphenes. The appearance of patterns of phosphenes allows otherwise blind patients to detect and recognize objects, orient themselves or navigate obstacles [1]. However, so far retinal implants have struggled to provide artificial vision useful in everyday life. There are multiple technological and physiological constraints that currently keep artificial vision far away from natural vision, among which are limitations on the electrode number and density, the size of the restored field of view, the unselective stimulation of retinal circuits or cells, and the temporal resolution of the elicited responses [2,3]. These limitations lead to exhaustion in users, who have to learn an almost totally new way to see, and are not able to count only on the implant for daily autonomous activities [4].

A key factor in artificial vision is the temporal resolution of phosphenes. The electrical stimulation of the retina with a fixed electrode causes the phosphene to fade in less than a second [5–7]: a phenomenon that makes continuous perception above flicker fusion nearly impossible. The fading of the phosphenes elicited by retinal implants presumably originates in the inner retinal circuits mediating contrast adaptation [6–8]. Similar to the adaptation process to static visual stimuli (Troxler effect), the engagement of these circuits during retinal ganglion cell (RGC) network-mediated stimulation by either epiretinal or subretinal implants generates an adaptive response to repeated static electrical stimuli [5,6,9]. The desensitisation rate increases with the stimulation frequency [7], and at the stimulation rates typically used for retinal stimulation, between 5 to 30 Hz, RGCs rapidly stop responding in less than a few hundredths of seconds. The physiological adaptation of the RGC response over repeated static stimulation observed in animal models is frequently associated with the phosphene fading reported by implanted patients during clinical trials [10,11]. This association is motivated by the similarity in the time courses of the two phenomena, characterised by a rapid drop followed by a slower decay [6,12,13].

In vision, ocular micromovements (e.g. microsaccades) refresh the image projected on the retina avoiding the Troxler effect [14]. In artificial vision, avoiding response adaptation is possible only when three conditions are met. First, the electrodes must be activated by natural or artificial light entering the eye through the pupil, such as for photosensitive prostheses (e.g. Alpha AMS [15], PRIMA [16] and POLYRETINA [17,18]), so that eye micromovements can effectively shift the stimulated area of the retina. Second, the stimulation must be at high spatial resolution. Microsaccades occur with a great variety of amplitudes (from 0.2 to 10 degrees) and frequencies (from 1 Hz to 20 Hz). Under stable vision conditions, most fixational microsaccades occur 1 to 2 times per second, and cover around 1 degree of visual angle [19]. The stimulation resolution of the array must be fine enough so that a shift of a few hundreds of micrometers (1 degree) is sufficient to move the projected image by one or two electrodes over the prosthesis leading to stimulation of different retinal areas. Third, physiological micromovements must be preserved, such as in age-related macular degeneration [16].

When electric based prostheses are used (e.g. Argus® II [11]), users must actively perform head movements to refresh the visual field viewed by the camera: a procedure learned during the first weeks of postoperative training. Alternatively, if eye movements are not dysfunctional and an eye tracker is used, large voluntary eye movements could be used to shift the region of interest within the camera’s field of view [20]. Although the patient’s performance typically improves during the first learning phase [21], it quickly reaches a plateau, and several patients described active scanning as an exhaustive task [4]. Recently, our group proposed a naturalistic spatiotemporal modulation strategy for electrical prostheses, which, in retinal explants, reduced retinal desensitisation and prolonged the RGC response upon static stimulation [7].

Simulated prosthetic vision (SPV) is a key tool to understand the minimal requirements of artificial vision [22–32]. However, so far, most SPV studies eluded the temporal properties of artificial vision, in particular phosphene fading. Only one study recently introduced fading using short video clips presented on a computer screen [33]. In virtual reality (VR), subjects are able to scan the environment using head movements, allowing the systematic investigation of how scanning influences fading. In this work, we introduced perceptual fading into a VR-based SPV, under the hypothesis that fading is solely caused by retinal desensitisation, according to previously reported temporal dynamics in RGCs [7]. We analysed the consequences of fading on a VR task with sighted participants, and simulated fading compensation approaches based on the previously described naturalistic spatiotemporal modulation strategies [7] to evaluate whether they could indeed successfully reduce participants’ head scanning.

## 2. METHODS

### 2.1 Ethical Statement

Experiments were approved by the human research ethics committee of École polytechnique fédérale de Lausanne (decision number 042–2018/16.10.2018). Ten sighted volunteers were involved in the study (**Table 1**).

**Table 1.**
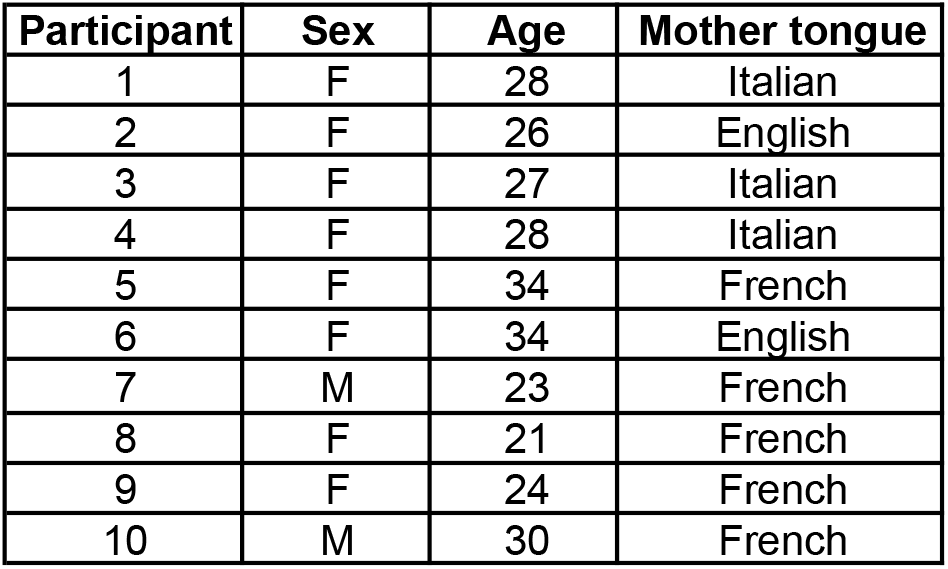
List of normally sighted volunteers enrolled in the study.

### 2.2 Prosthetic vision

SPV was generated by adapting a previously described approach [22]. The experiment was performed using a Dell Precision 3630 computer with an Intel Xeon E-2146G CPU (3.50 GHz) and an Nvidia GeForce GTX 1080 GPU. The VIVE Pro Eye head mounted display was used. Tracking of head orientation and position was provided using two VIVE Base Station 2.0 tracking cameras placed in opposite corners of the room (roughly 2.5 m apart) and aimed at the center of the room where the participant was sat. SPV was developed using Unity and computed using Cg shaders, allowing real-time operation. The code is available online (https://github.com/lne-lab/polyretina_vr). Both the spatial and temporal properties of phosphenes were considered. Images were converted into phosphenes distributed over a visual angle of 45 degrees using the layout of the POLYRETINA prosthesis (80/120, electrode diameter in μm / electrode pitch in μm). The total number of phosphenes was 9’914. For each frame, three sources of random variability were included: the phosphene size was randomly varied between ±30% of the electrode size, the phosphene brightness was randomly varied between 50% (grey) to 100% (white) of the phosphene’s default brightness, and 10% of the electrodes were considered not functional. Then, a distortion due to unintended activation of axon fibres was introduced (λ = 2) [22]. The frame rate of SPV was set to 5 Hz with a 11 ms frame duration (1 frame on / 17 frames off). SPV was presented to the right eye only.

### 2.3 Simulation of perceptual fading

A novel aspect of the SPV was the inclusion of perceptual fading, which is the reduction in brightness of phosphenes over a short amount of time due to retinal desensitisation. Concurrently, compensation approaches for fading were simulated based on the naturalistic spatiotemporal modulation strategies previously characterised with retinal explants from blind mice [7] (**Figure 1)**.

**Figure 1.**
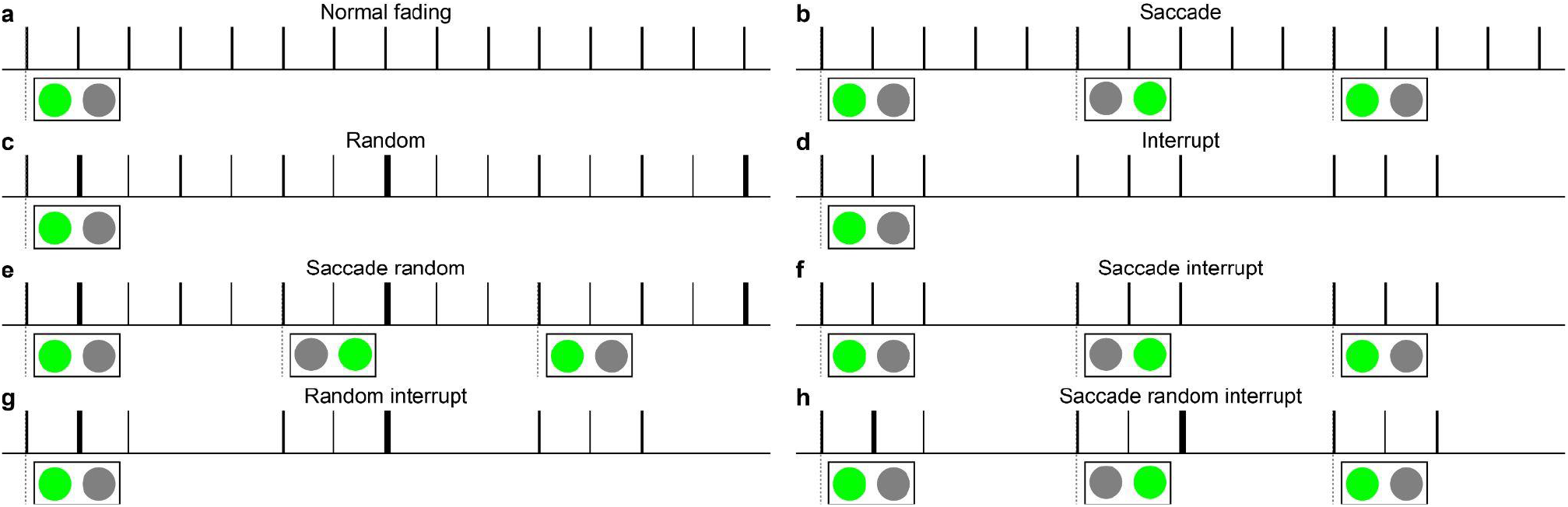
Sketches of the naturalistic spatiotemporal modulation strategy previously used in blind retinal explants to determine the decay in RGCs response and, consequently, the phosphene brightness [7].

In SPV, ‘normal fading’ (F) corresponds to the natural desensitisation of the retina obtained with 5 Hz repeated illumination of the same POLYRETINA electrode (**Figure 1a**), while ‘no fading’ (NF) corresponds to stable brightness over time. The ‘saccade’ strategy (S) is a spatial modulation strategy. It simulates natural microsaccades by having the entire image oscillate horizontally by the pitch of one phosphene once every second. This way, new phosphenes will become active and refresh at least some part of the prosthetic image, thus counteracting fading. The brightness decay was estimated from the response decay measured in RGCs by alternating the illumination of two neighboring electrodes at 1 Hz switching rate (**Figure 1b**). The ‘random’ (R) and the ‘interrupt’ (I) conditions are temporal modulation strategies. As before, the brightness decay was estimated from the response decay measured in RGCs. In the R condition, the duration of the light pulse was varied, leading to randomised inter-pulse durations (**Figure 1c**). In the I condition, for each second, only the first three out of five pulses were delivered (**Figure 1d).** The remaining conditions (**Figure 1e-h**), were a combination of the three strategies already described: saccade/random (SR, **Figure 1e**), saccade/interrupt (SI, **Figure 1f**), random/interrupt (RI, **Figure 1g**), and saccade/random/interrupt (SRI, **Figure 1h**). Last, the brightness reduction was approximated as a linear decay with two time constants (T_f_, fast decay constant; T_s_, slow decay constant) for each condition (**Table 2**).

**Table 2.** Fast and slow time constants for each condition were determined according to the persistence of RGC response observed in blind retinas upon stimulation with POLYRETINA [7].

| Condition | Decay Time (s) |  |  |  |
| --- | --- | --- | --- | --- |
|  | SPV |  |  | Experimental value from [7] |
| | $T_f$ | $T_s$ | Total | Total |
| F | 0.100 | 0.300 | 0.400 | 0.400 |
| S | 0.175 | 0.525 | 0.700 | 1.400 |
| R | 0.200 | 0.600 | 0.800 | 0.800 |
| I | 0.100 | 0.300 | 0.400 | 0.400 |
| SR | 0.175 | 0.525 | 0.700 | 1.400 |
| SI | 0.275 | 0.825 | 1.100 | 2.200 |
| RI | 0.800 | 2.400 | 3.200 | 3.200 |
| SRI | 0.550 | 1.650 | 2.200 | 4.400 |
| NF | - | - | - | - |

S strategy involves the alternation between two neighbouring electrodes. Therefore the experimental persistence of the RGC response was halved to account for the fading for each phosphene. Based on the time constants, the brightness of each phosphene was adjusted at each frame (**Table 3**).

**Table 3.** Brightness decay for one phosphene. For conditions involving S strategy (S, SI, SR and SRI), the table reports only the brightness of the first phosphene, since the brightness of the second phosphene depends on its activation history. Empty cells indicate that the frame was not displayed due to the I strategy.

| Time (s) | Percent phosphene brightness |  |  |  |  |  |  |  |  | Recovery |
| --- | --- | --- | --- | --- | --- | --- | --- | --- | --- | --- |
|  | F | S | R | I | SR | SI | RI | SRI | NF |  |
| 0.0 | 100.0 | 100.0 | 100.0 | 100.0 | 100.0 | 100.0 | 100.0 | 100.0 | 100.0 | 0.0 |
| 0.2 | 33.3 | 47.6 | 50.0 | 33.3 | 47.6 | 63.6 | 87.5 | 81.8 | 100.0 | 27.1 |
| 0.4 | 0.0 | 28.6 | 33.3 | 0.0 | 28.6 | 42.4 | 75 | 63.6 | 100.0 | 48.8 |
| 0.6 | 0.0 | 9.5 | 16.7 |  | 9.5 |  |  |  | 100.0 | 65.7 |
| 0.8 | 0.0 | 0.0 | 0.0 |  | 0.0 |  |  |  | 100.0 | 78.4 |
| 1.0 | 0.0 | 0.0 | 0.0 | 0.0 | 0.0 | 6.1 | 45.8 | 36.4 | 100.0 | 87.5 |
| 1.2 | 0.0 | 0.0 | 0.0 | 0.0 | 0.0 | 0.0 | 41.7 | 30.3 | 100.0 | 93.6 |
| 1.4 | 0.0 | 0.0 | 0.0 | 0.0 | 0.0 | 0.0 | 37.5 | 24.2 | 100.0 | 97.3 |
| 1.6 | 0.0 | 0.0 | 0.0 |  | 0.0 |  |  |  | 100.0 | 99.2 |
| 1.8 | 0.0 | 0.0 | 0.0 |  | 0.0 |  |  |  | 100.0 | 99.9 |
| 2.0 | 0.0 | 0.0 | 0.0 | 0.0 | 0.0 | 0.0 | 25.0 | 6.1 | 100.0 | 100.0 |
| 2.2 | 0.0 | 0.0 | 0.0 | 0.0 | 0.0 | 0.0 | 20.8 | 0.0 | 100.0 | 100.0 |
| 2.4 | 0.0 | 0.0 | 0.0 | 0.0 | 0.0 | 0.0 | 16.7 | 0.0 | 100.0 | 100.0 |
| 2.6 | 0.0 | 0.0 | 0.0 |  | 0.0 |  |  |  | 100.0 | 100.0 |
| 2.8 | 0.0 | 0.0 | 0.0 |  | 0.0 |  |  |  | 100.0 | 100.0 |
| 3.0 | 0.0 | 0.0 | 0.0 | 0.0 | 0.0 | 0.0 | 4.2 | 0.0 | 100.0 | 100.0 |
| 3.2 | 0.0 | 0.0 | 0.0 | 0.0 | 0.0 | 0.0 | 0.0 | 0.0 | 100.0 | 100.0 |

Brightness recovery followed an exponential growth according to classical models of neuronal excitability [34–36] and regardless of the strategy used (**Equation 1** and **Table 3**), where B is the phosphene brightness, t is the time the phosphene has been off, a is the time for the phosphene to completely recover (set to 2 s) and b defines the arc of recovery (set to 3).

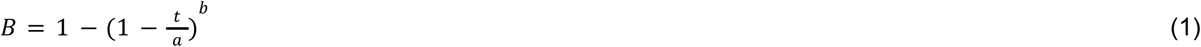

A representative example of image perception at various frames is shown in **Figure 2.**

**Figure 2.**
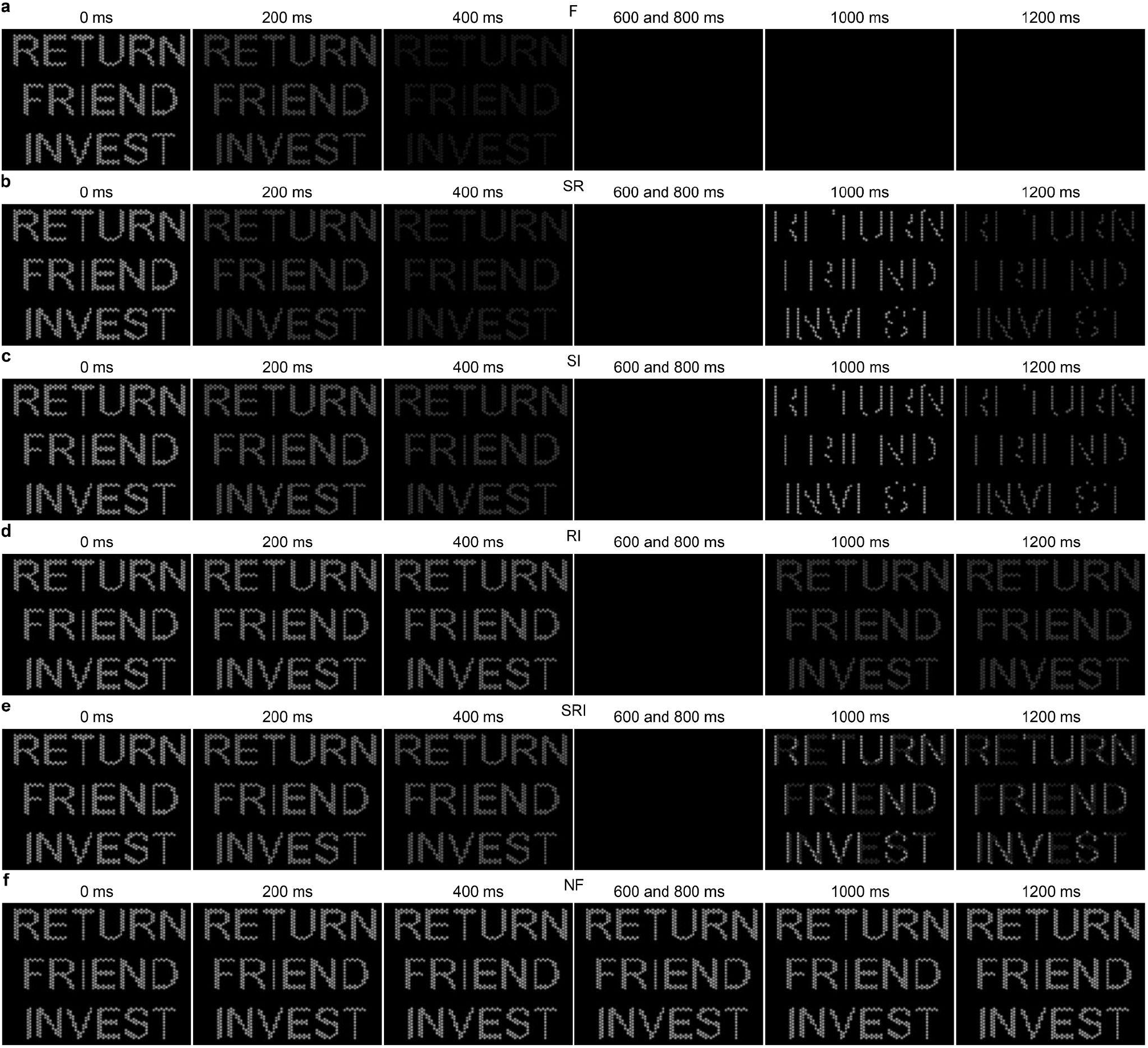
Example of perception with six strategies: F (**a**), SR (**b**), SI (**c**), RI (**d**), SRI (**e**) and NF (**f**).

The fading logic was applied simultaneously to each phosphene every time there was a stimulus presentation (**Figure 3**). The brightness of each phosphene is characterised by two values, T_a_ (the time since that phosphene’s first pulse) and T_n_ (the time since it’s most recent pulse). These values as well as a value keeping track of each phosphene’s potential brightness were updated in real-time on the GPU where the simulation is also being processed. To read/write such an amount of data on the GPU we used multiple render textures and accessed them in a double-buffered fashion to create a read/write data matrix where the values could be stored and updated in real-time. In other words, all processing and necessary data for the simulation remained solely on the GPU, where computations could be completed fast enough to enable real-time SPV.

**Figure 3.**
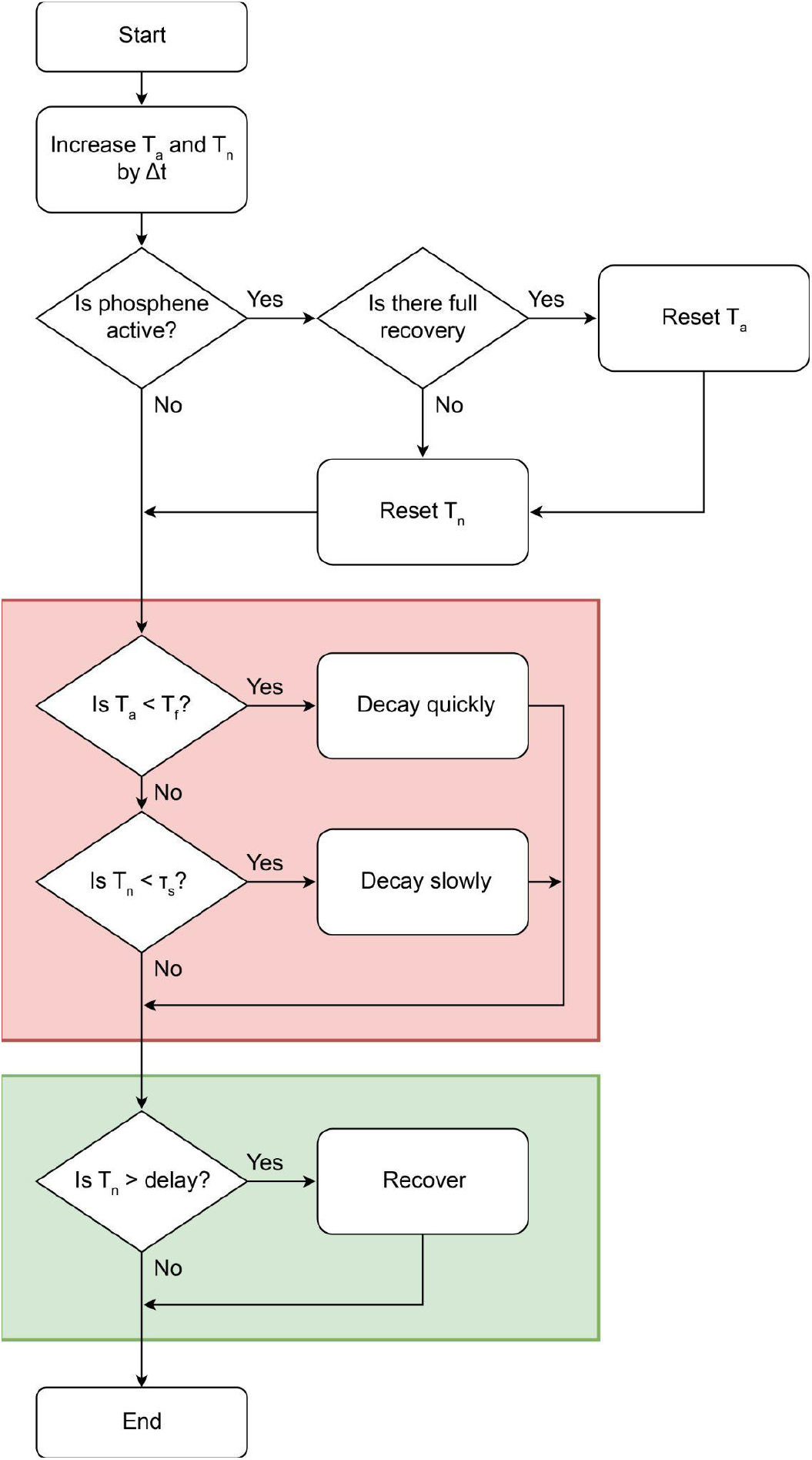
Fading logic. T_a_ is the time since the first pulse, T_n_ is the time since the last pulse, Δt is the time since the fading logic was last executed, T_f_ is the fast decay constant, T_s_ is the slow decay constant.

### 2.4 Experiment Design

The experiment consisted of a single repeated task in which participants had to read aloud three six-letter words. Words were presented in the mother tongue of the participant and on separate lines. Each word was 7 degrees tall and the entire stimulus (i.e., the three words) took up 27 degrees vertically and around 40 degrees horizontally. Participants could rotate their heads in order to adjust their view of the words, for example, if fading had completely hidden the words. Translational movements however did not affect the view of the stimulus. Once a participant had given their answers, the experimenter would mark the trial as finished and the participant would have a two second pause until the next trial. As participants were able to move their heads, their virtual head position was reset at the start of each trial.

The task was designed to not be too difficult but to also take time to complete. As the study was focused on the impact of phosphene fading, it was important that participants could not complete the task in such a short amount of time that the fading was not perceived. By having three words instead of one, it guaranteed that a participant could not complete the task before the image had completely faded prior to any accommodating head rotation. On the other hand, the words were displayed with a generous size, so that participants could actually complete it. SPV includes several laborious aspects of artificial vision such as a low spatial resolution, restricted field of view and refresh rate, as well as unintended visual distortions from axon fibre activation and perceptual desensitization. In pilot tests, making words smaller, in combination with the already unaccommodating SPV, increased the difficulty to unacceptable levels. Another important aspect of the task was that the three words fit entirely within the field of view of the prosthetic vision (45 degrees). By doing this, any head rotation from participants could be largely attributed to counteractive behaviour and not due to participants having to move their heads to look at the stimulus.

The task was completed under the 9 different conditions, presented randomly. Each condition was presented 5 times for a total of 45 trials per session. Participants were asked to come in for a total of six sessions to control for learning effects. Therefore, each participant completed 270 trials (30 per condition). Participants were given instructions on the task before starting. Most importantly, they were given the information that the SPV would fade over time and that head rotations could be used to counteract the fading.

### 2.5 Data analysis and plots

Data analysis was performed in MATLAB (MathWorks). Statistical analysis and graphical representation were performed with Prism 8 (Graph Pad). The D’Agostino & Pearson omnibus normality test was performed to justify the use of a non-parametric test. The box plots extend from the 25th to 75th percentiles. The line is the median. The + is the mean. The whiskers extend to the smallest and the largest value. In plots, p-values were reported as: * p < 0.05, ** p < 0.01, *** p < 0.001, and **** p < 0.0001.

## 3. RESULTS

We collected two indicators for each trial: the time to complete the task and the head rotation.

### 3.1 Time to complete the task

First, we evaluated the time taken by each subject to complete the task (**Figure 4**). This parameter provides the first information on the performance for each condition.

**Figure 4.**
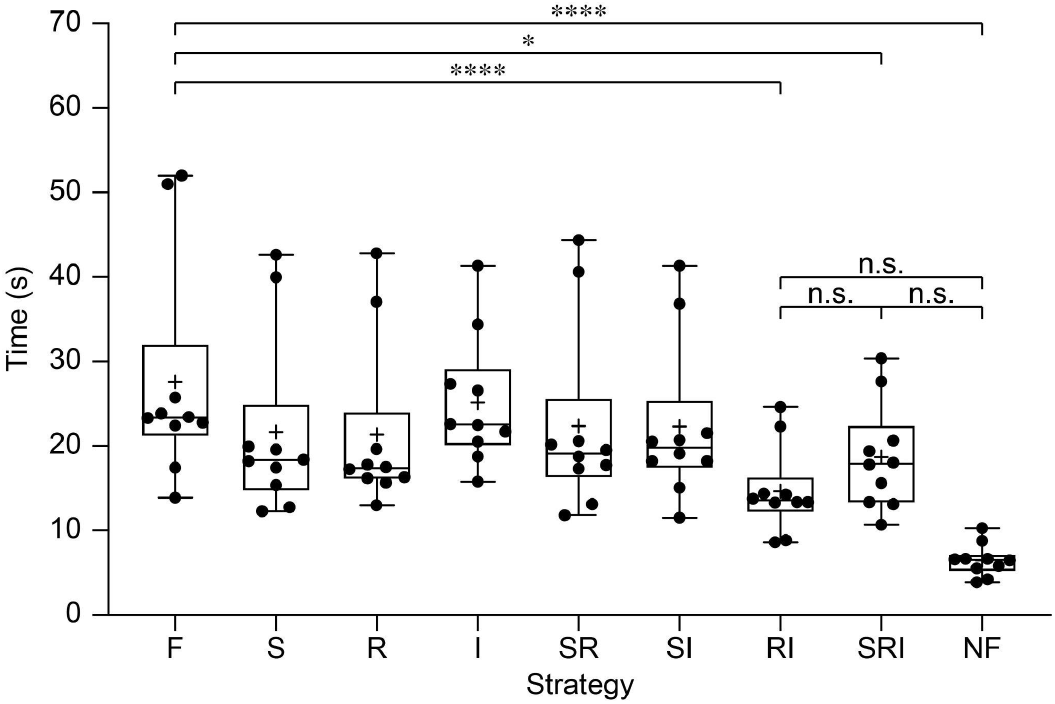
Quantification of the time taken by each subject (mean ± s.d.). For each condition, each subject performed 30 trials, and the time was averaged among trials. Individual data points are the ten subjects.

The Friedman one-way repeated measure analysis of variance by ranks reported a significant difference among the conditions (p < 0.0001). Dunn’s multiple comparison test showed that three conditions resulted in a time to complete the task significantly lower than the fading condition: RI (p < 0.0001), SRI (p = 0.0393) and NF (p < 0.001). The remaining conditions (S, R, I, SR, SI) do not result in a statistically significant difference in the time to complete the task compared to the fading condition (**Table 4**), indicating that those strategies might not be as powerful as the others. Interestingly, the RI and SRI strategies are not only significantly better than F condition, but also reported results statistically similar to the NF condition, which indicates that they are extremely efficient in counteracting fading. The other strategies (S, R, I, SR, SI) are significantly worse than NF condition, which confirms their poor performance in counteracting fading. In summary, the most promising strategies to counteract fading were RI and SRI.

**Table 4.**
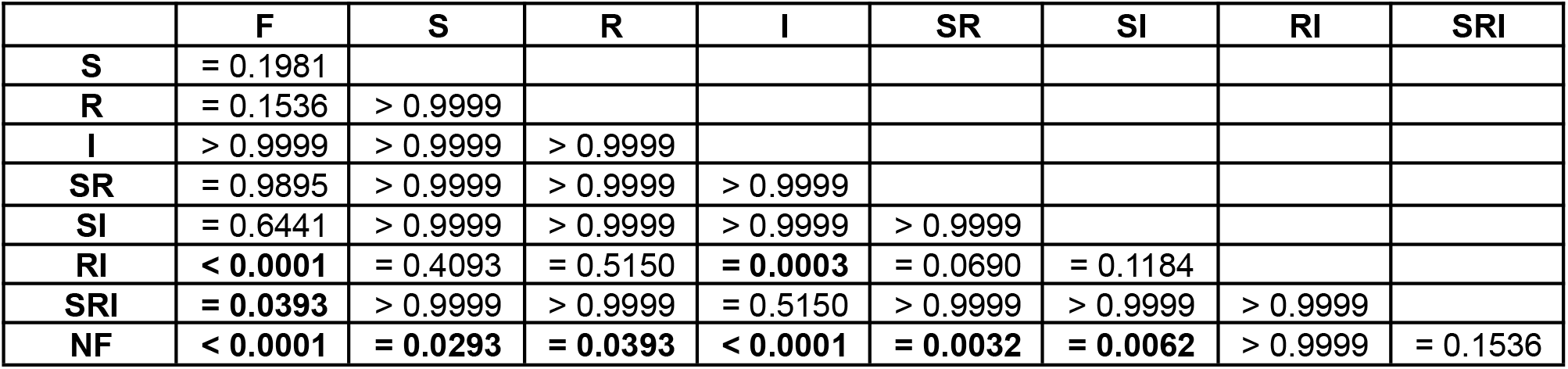
p-values from the Dunn’s multiple comparison test. Significant comparisons are in bold (p < 0.05).

|  | F | S | R | I | SR | SI | RI | SRI |
| --- | --- | --- | --- | --- | --- | --- | --- | --- |
| S | = 0.1981 |  |  |  |  |  |  |  |
| R | = 0.1536 | > 0.9999 |  |  |  |  |  |  |
| I | > 0.9999 | > 0.9999 | > 0.9999 |  |  |  |  |  |
| SR | = 0.9895 | > 0.9999 | > 0.9999 | > 0.9999 |  |  |  |  |
| SI | = 0.6441 | > 0.9999 | > 0.9999 | > 0.9999 | > 0.9999 |  |  |  |
| RI | < 0.0001 | = 0.4093 | = 0.5150 | = 0.0003 | = 0.0690 | = 0.1184 |  |  |
| SRI | = 0.0393 | > 0.9999 | > 0.9999 | = 0.5150 | > 0.9999 | > 0.9999 | > 0.9999 |  |
| NF | < 0.0001 | = 0.0293 | = 0.0393 | < 0.0001 | = 0.0032 | = 0.0062 | > 0.9999 | = 0.1536 |

### 3.2 Head Rotation

In order to counteract the fading of phosphenes during prosthetic vision, implanted patients are instructed to continuously move their head. The side-effect is a decline in comfort of the use of the device from constant scanning. We hypothesised that head movements could be reduced by employing strategies to slow down the fading.

The head rotation data consisted of a non-zero amount of quaternion values depending on the duration of the trial (**Figure 5a)**. To analyse the data, each quaternion was first converted into rotations around the vertical and lateral axes (**Figure 5b**), since rotations were almost absent in the longitudinal axis. Then, head rotation data was split into two separate measurements: one focused on the magnitude of head rotations and the other on the spread of head rotations.

**Figure 5.**
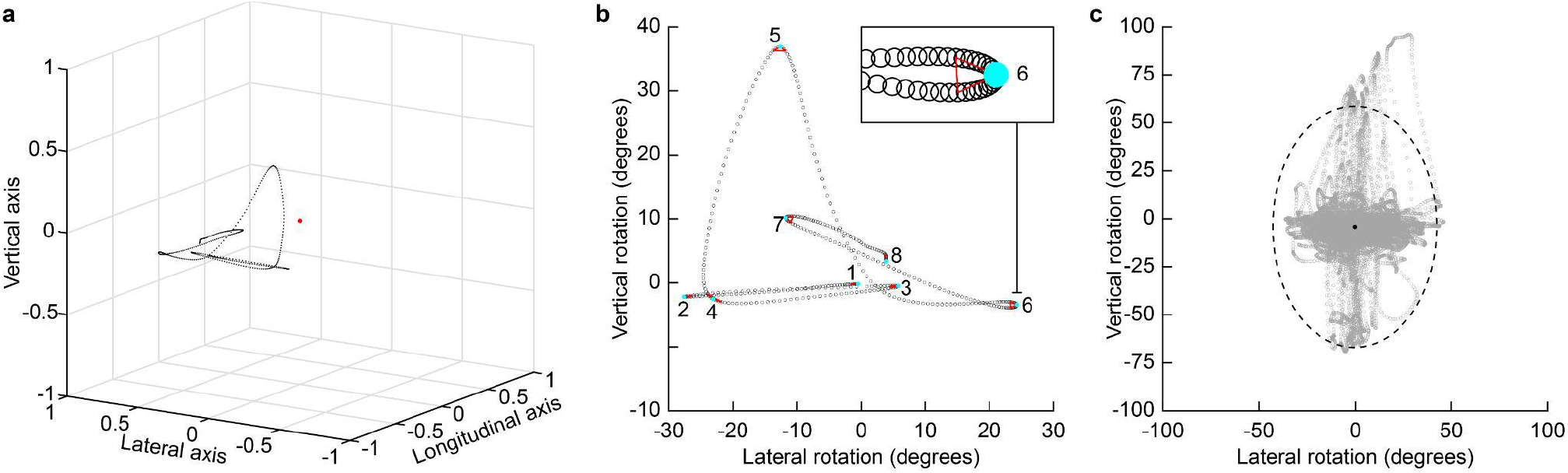
Representative example of head rotation data in one trial under condition F. (**a**) Example of the quaternion data. The red dot is the position of the head, while the black dots are the coordinates at which the participant was looking throughout the trial. The vertical axis is the yaw, the lateral axis is the pitch and the longitudinal axis is the roll. (**b**) Two dimensional plot of the head rotations along the lateral and vertical axes. Cyan circles are the peaks at the extremities of the segments, while the red triangles represent the angle of the change in head rotation at the peak. The insert shows a magnified view of a detected peak. (**c**) Spread of the head rotations along the lateral and vertical axes. The black dashed line is three standard deviations of the data for both axes. While the data is not normally distributed, Chebyshev’s theory of inequality states that at least 88.89% of values will lie within the ellipsoid. The black circle is the mean head rotation.

To measure the magnitude, head rotations from each trial were segmented into individual rotation vectors (**Figure 5b**). Segments were identified using a peak analysis on the head’s inverse velocity. Segments were then filtered using the k-means algorithm using the peak’s height, width and the turning angle at the point of the peak. The first and last data points of each trial were always marked as peaks. If multiple data points were identified for a single segment then the data point closest to the correct k-means cluster centre was chosen. The segment magnitude was calculated by summing the rotation between each data point in the segment. Finally, the magnitude of these vectors was averaged. This approach was chosen over simpler options such as the total head rotation as this was heavily affected by the time taken to complete the task. Similarly, head velocity proved to be a misleading metric as participants’ head rotations were often discontinuous, pausing while participants focused on the stimulus and then subsequently moving as fading began to degrade vision. As a consequence of the stop-start nature of head rotations the data could be easily segmented into individual rotation vectors, with an associated magnitude.

To measure the spread of head rotations (**Figure 5c**), we calculated the average rotation of the head away from the stimulus for each trial, using the dist function in MATLAB. This function returns the angle between two quaternions in unit radians. Using this function in combination with an identity quaternion angled directly towards the stimulus, an angle was calculated for each head rotation which was centered around the stimulus. Finally, the data was converted into degrees from radians and averaged per trial.

#### 3.2.1 Magnitude of head rotations

During trials under normal fading (F), we observed large head rotations away from the stimulus (**Figure 6a,b**), while head rotations were virtually absent during the NF condition (**Figure 6e,f**). Spatiotemporal modulation strategies allowed the reduction of head rotations by a variable factor depending on the strategy (**Figure 6c,d** for RI strategy).

**Figure 6.**
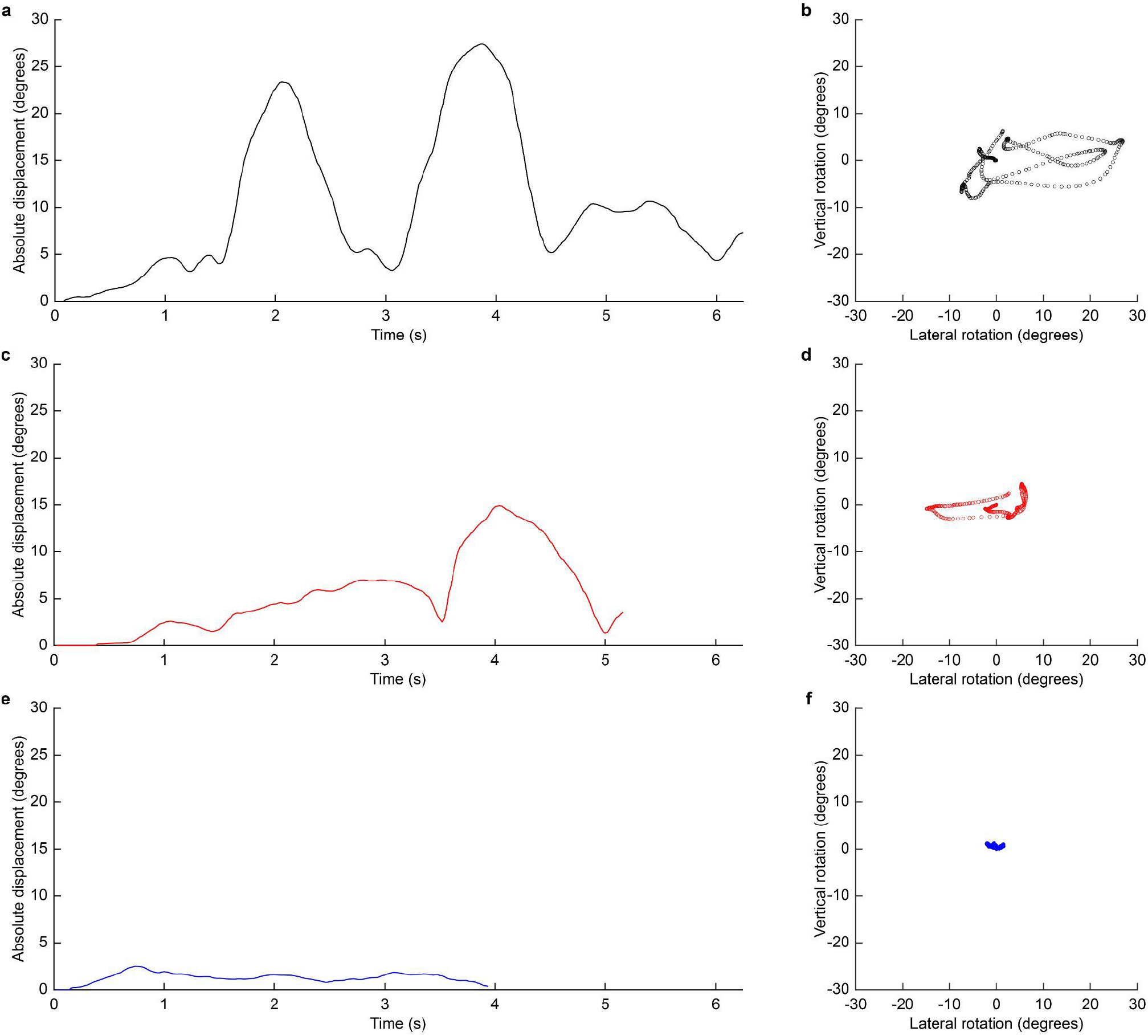
(**a**, **c**, **e**) Representative example of absolute angular rotation of the participant’s head from the stimulus over time for conditions F (**a**), RI (**c**) and NF (**e**) during one trial in the same session. (**b**, **d**, **f**) Corresponding vertical and lateral rotations for conditions F (**b**), RI (**d**) and NF (**f**) during a single trial.

The average magnitude of the head rotations was calculated then for all the conditions to determine which strategy would provide a statistically significant reduction (**Figure 7**).

**Figure 7.**
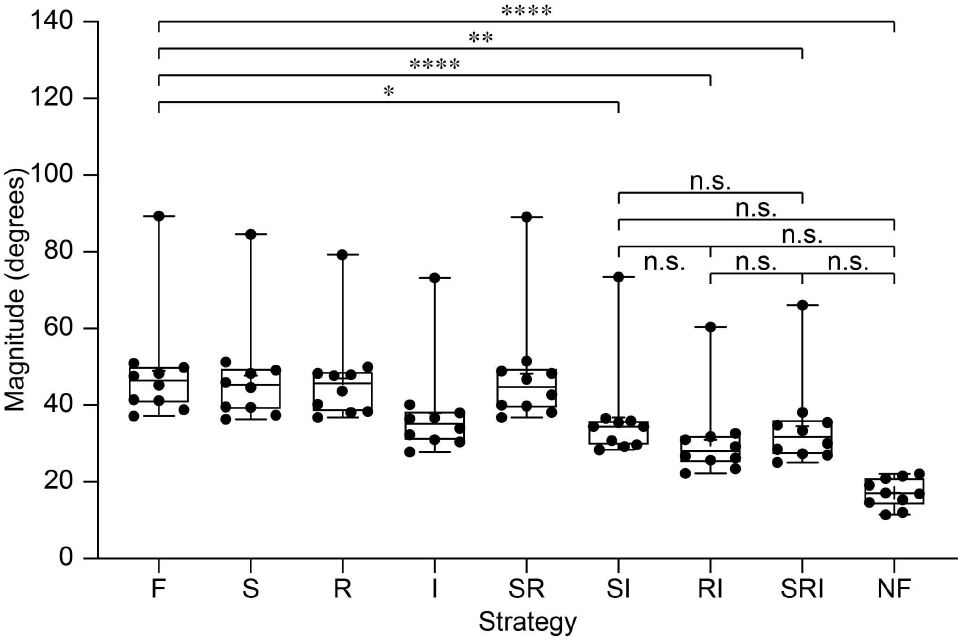
Quantification of the magnitude of head rotations per subject (mean ± s.d.). For each condition, each subject performed 30 trials, and data was averaged among trials. Individual data points are the ten subjects.

The Friedman one-way repeated measure analysis of variance by ranks reported a significant difference among the conditions (p < 0.0001). Similar to the time to complete the task, Dunn’s multiple comparison test showed that four conditions resulted in a magnitude of head rotations significantly lower than the fading condition: SI (p = 0.0293), RI (p < 0.0001), SRI (p = 0.0023) and NF (p < 0.001). The remaining conditions (S, R, I, SR) do not result in a significant difference in the magnitude of head rotations compared to the fading condition (**Table 5**), indicating that those strategies might not be as powerful as the others. Interestingly, SI, RI and SRI strategies are not only significantly better than the F condition, but also reported results statistically similar to the NF condition, which indicates that they are extremely efficient in counteracting fading and reducing the magnitude of head rotations. The other strategies (S, R, I, SR) are significantly worse than the NF condition, which confirms their poor performance. In summary, the most promising strategies to reduce the magnitude of head rotations were SI, RI and SRI.

**Table 5.**
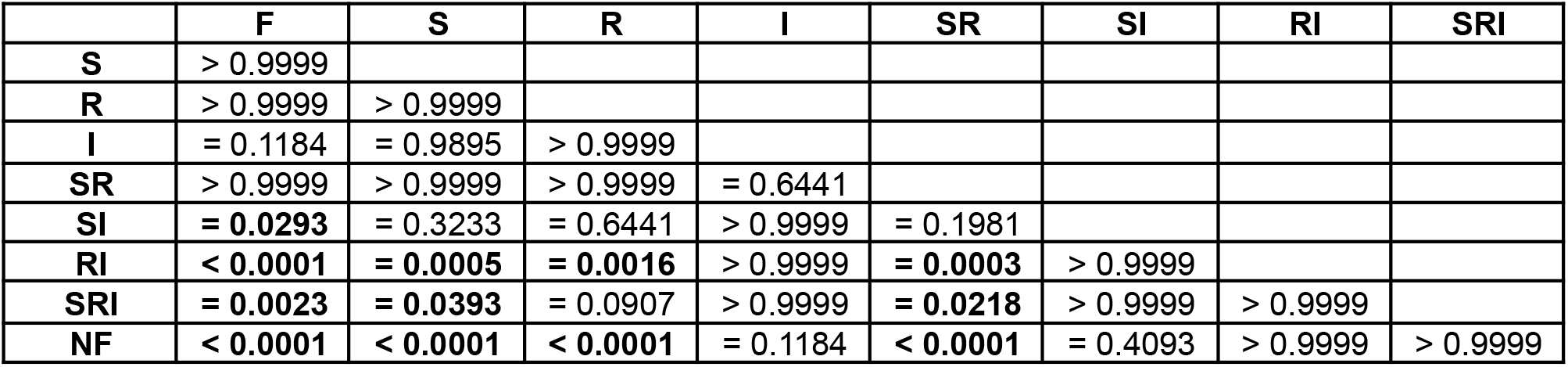
p-values from the Dunn’s multiple comparison test. Significant comparisons are in bold (p < 0.05).

#### 3.2.2 Spread of head rotations

Next, we quantified the spread of the head rotations. Similar to the magnitude, during trials under normal fading (F), we observed a spread of rotations away from the stimulus in both vertical and lateral direction (**Figure 8**, black). Modulation strategies reduce the spread by a variable factor depending on the strategy (**Figure 8**, red for RI strategy), which are further reduced in NF condition (**Figure 8**, blue).

**Figure 8.**
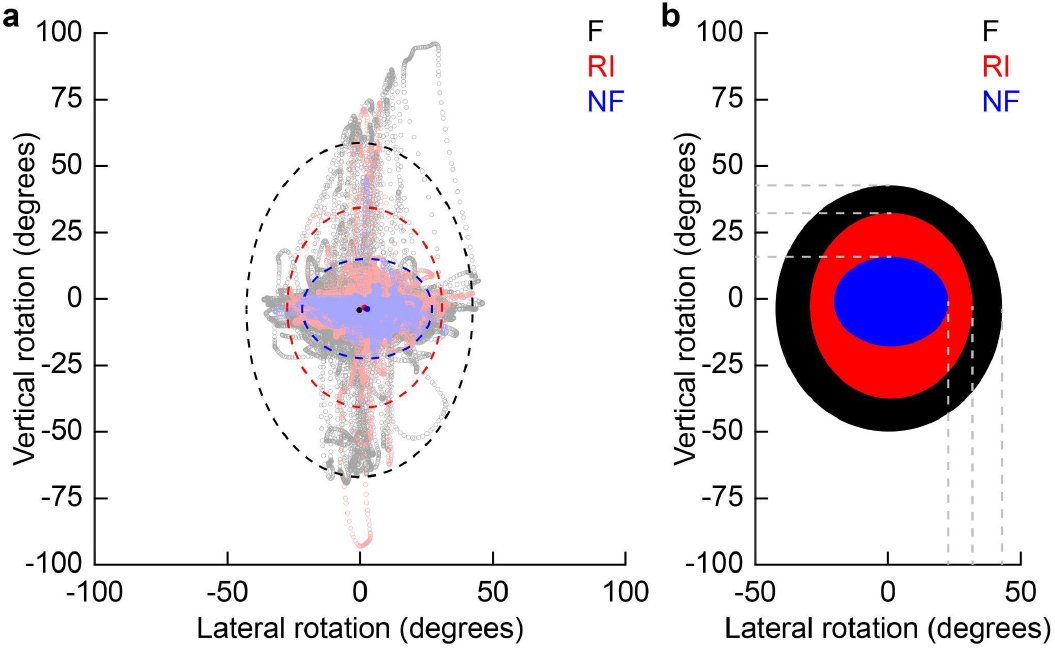
(**a**) Representative plot of the head rotations from one participant under F (black), RI (red) and NF (blue) conditions. The circles in the centre represent the mean of the data and the dashed ellipsoids indicate three standard deviations of the data for both axes. (**b**) Ellipsoids represent the average across all participants of the three standard deviations of the data distributions.

The average spread of the head rotations was calculated then for all the conditions to determine which strategy would provide a statistically significant reduction (**Figure 9**).

**Figure 9.**
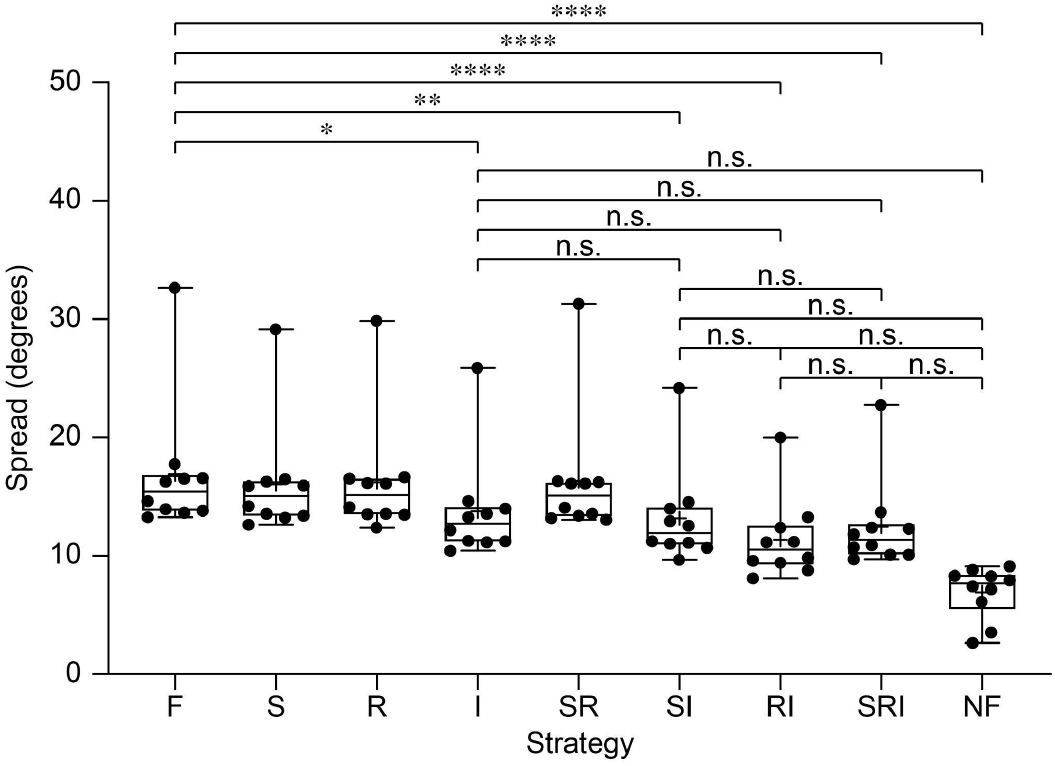
Quantification of the spread of head rotations per subject (mean ± s.d.). For each condition, each subject performed 30 trials, and data was averaged among trials and subjects. Individual data points are the ten subjects.

The Friedman one-way repeated measure analysis of variance by ranks reported a significant difference among the conditions (p < 0.0001). Similar to the previous analysis, Dunn’s multiple comparison test showed that five conditions resulted in a spread of head rotations significantly lower than the fading condition: I (p = 0.0393), SI (p = 0.0016), RI (p < 0.0001), SRI (p < 0.0001) and NF (p < 0.001). Also, the remaining conditions (S, R, SR) do not result in a significant difference in the spread of head rotations compared to the fading condition (**Table 6**), indicating that those strategies might not be as powerful as the others. Interestingly, I, SI, RI and SRI strategies are not only significantly better than the F condition, but also reported results statistically similar to the NF condition, which indicate that they are extremely efficient in counteracting fading and reducing the spread of head rotations. The other strategies (S, R, SR) are significantly worse than the NF condition, which confirms their poor performance. In summary, the most promising strategies to reduce the spread of head rotations were I, SI, RI and SRI.

**Table 6.** p-values from the Dunn’s multiple comparison test. Significant comparisons are in bold (p < 0.05).

|  | F | S | R | I | SR | SI | RI | SRI |
| --- | --- | --- | --- | --- | --- | --- | --- | --- |
| S | > 0.9999 |  |  |  |  |  |  |  |
| R | > 0.9999 | > 0.9999 |  |  |  |  |  |  |
| I | **= 0.0393** | > 0.9999 | > 0.9999 |  |  |  |  |  |
| SR | > 0.9999 | > 0.9999 | > 0.9999 | > 0.9999 |  |  |  |  |
| SI | **= 0.0016** | = 0.6441 | = 0.1536 | > 0.9999 | = 0.5150 |  |  |  |
| RI | **< 0.0001** | **= 0.0045** | **= 0.0005** | = 0.8008 | **= 0.0032** | > 0.9999 |  |  |
| SRI | **< 0.0001** | = 0.0907 | = 0.0161 | > 0.9999 | = 0.0690 | > 0.9999 | > 0.9999 |  |
| NF | **< 0.0001** | **< 0.0001** | **< 0.0001** | = 0.0522 | **< 0.0001** | = 0.6441 | > 0.9999 | > 0.9999 |

### 3.3 Evaluation of the learning effect

Last, we asked if there was any learning effect during the task. Participants completed six sessions in total, spread out over at most two weeks, in order to see what effect practice would have on performance. The dataset was then separated for each session (**Figure 10**). For the time to complete the task, we observed a marked learning effect, shown by a performance increase after each session (**Figure 10a**). When only the last sessions for each conditions are compared, all conditions approach a qualitatively similar performance not significantly different than the F condition (p > 0.9999 for conditions S, R, I, SR, SI, RI and SRI; Kruskal-Wallis with Dunn’s multiple comparison test), except for the NF condition which remains significantly better than the F condition (p = 0.0006; Kruskal-Wallis with Dunn’s multiple comparison test).

**Figure 10.**
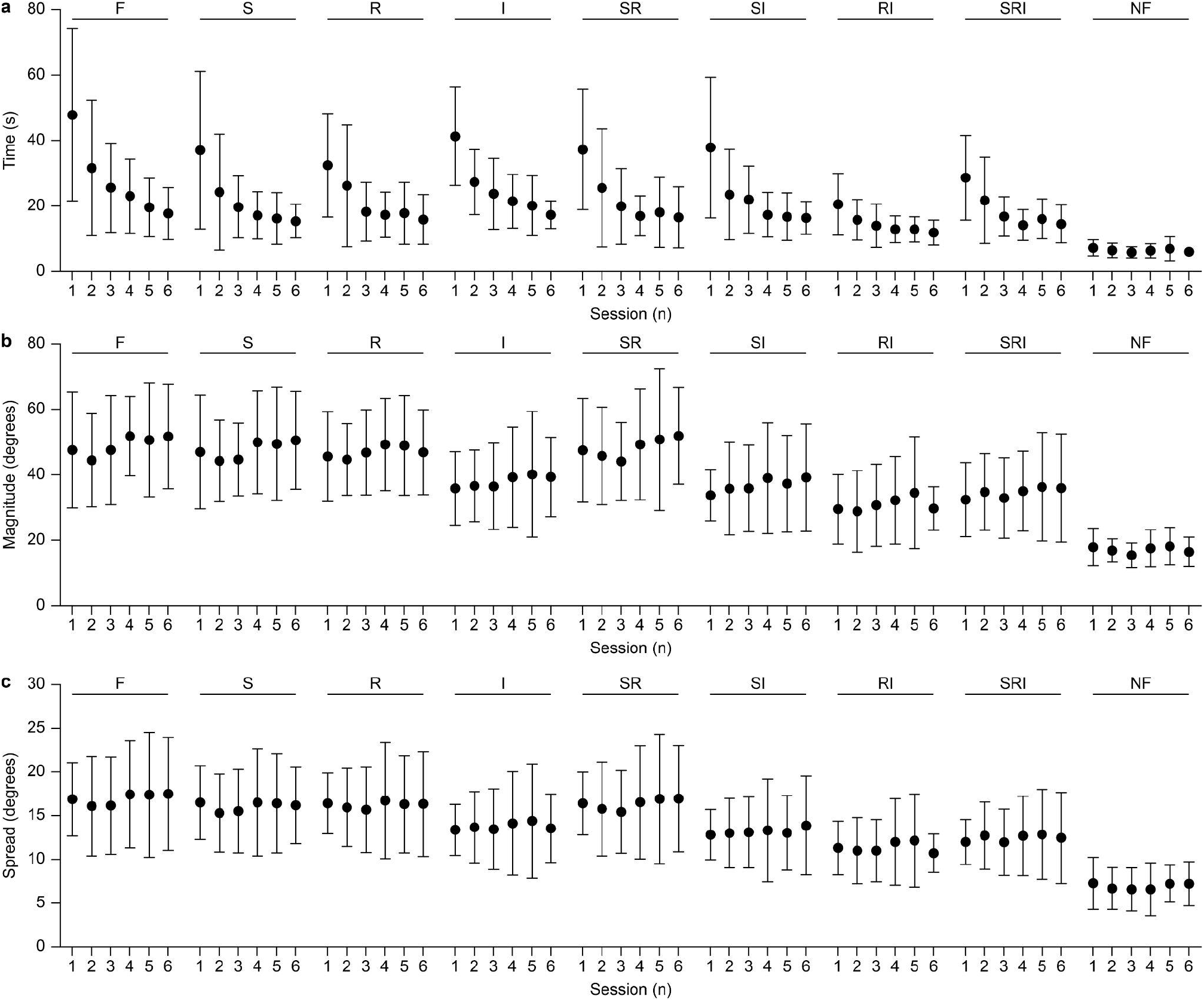
Quantification (mean ± s.d.) of the learning effect on the time taken to complete the task (**a**), the magnitude of head rotations (**b**) and the spread of head rotations (**c**). For each condition, each subject performed 30 trials, and data was averaged among trials and subjects.

This result indicates that just learning helps subjects to reduce the time to complete the task, independently of the use of fading compensation strategies. However, the situation is remarkably different for head rotations. Learning is not observed for both the magnitude (**Figure 10b**) and the spread (**Figure 10c**) of head rotations. For the magnitude, considering only the last session for each condition, RI strategy is significantly better than the F condition (p = 0.0049; Kruskal-Wallis with Dunn’s multiple comparison test) together with NF (p < 0.0001; Kruskal-Wallis with Dunn’s multiple comparison test); also RI and NF showed a statistically similar magnitude of head rotations (p > 0.9999; Kruskal-Wallis with Dunn’s multiple comparison test). A similar result was shown for the spread of head rotations. When considering only the last session for each condition, RI strategy is significantly better than the F condition (p = 0.0084; Kruskal-Wallis with Dunn’s multiple comparison test) together with NF (p < 0.0001; Kruskal-Wallis with Dunn’s multiple comparison test); also RI and NF showed a statistically similar magnitude of head rotations (p > 0.9999; Kruskal-Wallis with Dunn’s multiple comparison test).

In summary, this data indicates that practice improves the time to complete the task, but only modulation strategies (in particular the RI strategy) can significantly reduce the burden associated with active head scanning.

## 4. DISCUSSION

The perceptual fading of phosphenes limits the temporal resolution of artificial vision. The most naturalistic way to overcome fading is by exploiting spontaneous ocular micromovements, however this is possible only for high-resolution implants, activated by light and when physiological micromovements are preserved. Patients implanted with the PRIMA devices experienced flicker fusion [16] as these three conditions were met. However, this ideal situation might not be reached depending on the implant type and the patient condition. The clinical trial of Alpha IMS showed that the usefulness of natural eye movements for artificial vision varied among patients. Some patients could perceive complex objects like grating patterns or letters almost continuously. Eye tracking showed the presence of square waves, small amplitude drifts and microsaccades during phosphene perception [37], shifting the input image by 1 to 3 electrodes [38]. However, severe visual impairment is frequently associated with aberrant oculomotor functions, such as nystagmus and reduced saccade amplitude [39], which ultimately affects prosthetic vision. Therefore, other patients could not experience stable perception during retinal stimulation. Lastly, artificial vision in patients implanted with devices that cannot exploit eye movements (if preserved) will always be affected by perceptual fading.

As such, there is a wide interest in developing compensatory strategies to reduce, or ideally avoid, fading in artificial vision. It was previously shown that the randomization of the pulse duration lengthens the response persistence in RGCs [40] and electrically-evoked potentials in V1 [41]. Capitalising on these results, we proposed additional strategies to compensate for fading, such as interrupted train sequences and alternation between neighbouring electrodes to mimic artificial saccades [7] and showed that these strategies and their combination could lengthen the response of RGCs to electrical stimulation in explanted blind retinas.

In this article, we tested in SPV the effect of these strategies and their combination to reduce the voluntary head rotation imposed by the fading of artificial vision. The results showed that naturalistic modulation strategies help to reduce the magnitude and spread of head rotation during artificial vision. The use of an interrupted sequence appeared to be the most efficient strategy, in particular when combined with the randomisation of the stimulus duration. The intuition behind the interrupted sequence is to skip stimuli which will not lead anyway to any elicited phosphenes due to fading. For example, we measured in explanted retinas that for a 5 Hz repetition rate, only the first three stimuli will elicit a RGC response, therefore the next two stimuli will be skipped to allow response recovery. Next, we evaluated the possible effect of learning. Although we found an improvement in the time to complete the task, we did not observe a learning effect in head rotation. This result implies that active scanning due to fading is not expected to significantly improve with experience. Instead, an RI stimulation protocol can reduce the burden of active scanning by limiting the magnitude and spread of head rotations.

Contrary to the previous study in blind retinas [7], in SPV the implementation of a spatial modulation strategy (S) did not have a strong contribution on head rotations. The stimulation with retinal explants was limited to the alternation of two neighbouring POLYRETINA electrodes. Therefore, SPV was implemented by having the entire image oscillate horizontally by the pitch of one phosphene once every second. This choice was dictated by the nature of our previous experimental results from which time constants were derived. It should not be excluded that a more complex naturalistic modelling of ocular micromovements, involving random oscillations of the image in multiple directions might provide better results.

In conclusion, this study shows the relevance of stimulation strategies which could counteract perceptual fading to reduce active head scanning during prosthetic vision. Also, while this study was specifically designed for retinal implants, the results and their implications could be translated also to other visual implants, like optic nerve or cortical devices [42,43].

## 5. ACKNOWLEDGEMENT

This work was supported by École polytechnique fédérale de Lausanne, Medtronic plc, the foundation E. et G. Gelbert, and the European Union’s Horizon 2020 research and innovation programme under the Marie Skłodowska-Curie grant agreement No 861423.

J.T.T. designed the experiment, wrote the code and ran the tests. N.A.L.C. designed the experiment. S.H. ran the tests. M.C. ran the tests. D.G. designed and led the study, and wrote the manuscript. All the authors read and accepted the manuscript.

The authors declare no competing financial interests.

